# Kv7 channel antagonists block glycine receptors

**DOI:** 10.1101/2022.03.02.482705

**Authors:** Hsin-Wei Lu, Gabriel E. Romero, Pierre F. Apostolides, Hai Huang, Laurence O. Trussell

**Affiliations:** Neuroscience Graduate Program, Oregon Health and Science University, Portland OR 97239, USA; Physiology & Pharmacology Graduate Program, Oregon Health and Science University, Portland OR 97239, USA; Oregon Hearing Research Center & Vollum Institute, Oregon Health and Science University, Portland OR 97239, USA; Laboratory of Auditory Neurophysiology, Katholiecke Universität, Leuven, Belgium; Dept. of Neurobiology, Harvard Medical School, Boston, MA, USA; Kresge Hearing Research Institute and Dept. of Otolaryngology, University of Michigan Medical School, Ann Arbor, MI, USA; Dept. of Cell and Molecular Biology, Tulane University, New Orleans, LA, USA

## Abstract

XE991 (10,10-bis(4-pyridinylmethyl)-9(10H)-anthracenone) is currently the most widely used and specific antagonist of the Kv7 (KCNQ) family of K^+^ channels. We report an unexpected antagonistic effect of this drug on ionotropic glycine receptors. In recordings of synaptic transmission in two brainstem nuclei (the medial nucleus of the trapezoid body and the dorsal cochlear nucleus), 10 μM XE991, a concentration typical for Kv7 studies in brain tissue, inhibited evoked glycinergic inhibitory postsynaptic currents (IPSCs) without altering paired-pulse ratio, and also reduced the amplitude of glycinergic miniature IPSCs. These results are indicative of a direct effect of the drug on postsynaptic glycine receptors. XE991 also produced dose-dependent block of the response to exogenously applied glycine, to a degree comparable to the block of synaptic transmission. Moreover, the drug inhibited homomeric glycine receptors expressed on presynaptic membrane of the calyx of Held. The degree of block was independent of glycine concentration, suggesting an allosteric interaction. The effects of XE991 on glycine responses are not likely to reflect block of the glycine-activated Cl^-^ channels themselves, because block was voltage independent, and because GABA-activated Cl^-^ currents were resistant to XE991 at concentrations up to 100 µM. Linopirdine, but not retigabine, also antagonized glycine receptor currents. Given the prevalence of glycine receptor signaling in the brain, these observations should be taken into account in studies of the roles of Kv7 channels in neural circuit function and disease.

## Introduction

Kv7 (KCNQ) channels comprise 5 subtypes and play essential roles in neuronal circuits of the brain and sensory transduction, as well as cardiac and smooth muscle (Jespersen *et al*., 2005; Brown and Passmore, 2009; Haick and Byron, 2016). Accordingly, much effort has been made in developing a robust pharmacology for these channels, resulting in an extensive panel of specific activators and inhibitors of the channels. XE991 and linopirdine are the most commonly used antagonists, with low micromolar efficacy at blocking currents generated by native and heterologously expressed Kv7 channels (Grunnet *et al*., 2014). The value of such antagonists is realized in studies of the physiological roles of Kv7 channels and the potential development of novel treatments. Indeed, linopirdine, and later its derivative XE991, has been explored for cognitive enhancement in Alzheimer’s disease (Zaczek *et al*., 1998). At a cellular level, these compounds have been shown to induce hyperexcitability of cortical and hippocampal axons (Shah *et al*., 2008; Battefeld *et al*., 2014), enhance excitatory transmitter release in the auditory system (Huang and Trussell, 2011), and induce contraction of pulmonary arteries (Haick and Byron, 2016), all suggesting key roles of these channels in stabilizing membrane potential.

Based on the effects previously observed on excitatory neurotransmitter release, we examined the effects of XE991 on glycinergic inhibitory transmission in two brain regions, the medial nucleus of the trapezoid body (MNTB) of rat and mouse, and the dorsal cochlear nucleus (DCN) of mouse. These regions were chosen because of the dominant role of glycine in auditory brainstem function (Borst and Soria van Hoeve, 2012; Trussell and Oertel, 2018). We also examined both postsynaptic (heteromeric) and presynaptic (homomeric) alpha1 glycine receptors. We found that synaptic transmission mediated by glycine receptors was markedly inhibited by doses of XE991 typically used to block Kv7 channels. This effect was accounted for by an antagonism of glycine receptors, and was specific in that GABA_A_ receptors, which gate a similar Cl^-^ channel as glycine, were resistant to the drug. Similar effects were seen with presynaptic, homomeric glycine receptors, indicating that XE991 binds the alpha1 subunit of the receptor. Given the overlap in the pharmacology of native GABA and glycine receptors (Jonas *et al*., 1998; Beato *et al*., 2007; Li and Slaughter, 2007), XE991 is thus far unusual in being a reversible low-affinity glycine receptor antagonist with little activity at GABA_A_ receptors. However, this activity complicates the interpretation of studies of Kv7 channels, particularly in vivo.

## Methods

Experiments were performed using brain slices taken from P8-12 Wistar rats (for MNTB), and P8-22 (for MNTB) and P15-24 (for DCN) C57BL6 mice, according to protocols approved by the OHSU IACUC. Slices were prepared using a Leica VT1200S vibratome. For slices of rat or mouse MNTB, 180–220 μm thick coronal sections were prepared in an ice-cold solution containing (in mM) 230 sucrose, 25 glucose, 2.5 KCl, 3 MgCl_2_, 0.1 CaCl_2_, 1.25 NaH_2_PO_4_, 25 NaHCO_3_, 0.4 ascorbic acid, 3 myo-inositol, and 2 Na-pyruvate, bubbled with 5% CO_2_/95% O_2_. Immediately after cutting, slices were incubated at 35°C for 30–60 min in normal artificial cerebrospinal fluid (ACSF) and thereafter stored at room temperature. The ACSF for incubation and recording contained (in mM) 125 NaCl, 25 glucose, 2.5 KCl, 1 MgCl_2_, 2 CaCl_2_, 1.25 NaH_2_PO_4_, 25 NaHCO_3_, 0.4 ascorbic acid, 3 myo-inositol, and 2 Na-pyruvate, pH 7.4 bubbled with 5% CO_2_/95% O_2_.

For mouse DCN slices, 210– 230 μm coronal brainstem slices were cut in ice-cold solution containing the following (in mM): 87 NaCl, 25 NaHCO_3_, 25 glucose, 75 sucrose, 2.5 KCl, 1.25 NaH_2_PO_4_, 0.5 CaCl_2_, and 7 MgCl_2_, and bubbled with 5% CO2/95% O2. After cutting, slices were allowed to recover at 34°C in an artificial CSF (ACSF) solution containing the following (in mM): 130 NaCl, 2.1 KCl, 1.7 CaCl_2_, 1 MgSO_4_, 1.2 KH_2_PO_4_, 20 NaHCO_3_, 3 Na-HEPES, and 10–12 glucose, bubbled with 5% CO_2_/95% O_2_ (300–310 mOsm). After a 30–45 min recovery period, slices were kept at room temperature (22°C) until recording.

### Whole-Cell Recordings

All recordings were made at 33-34°C. Slices were transferred to a recording chamber and were continually perfused with ACSF (2–3 ml/min) at room temperature, except as noted. Neurons were viewed using Dodt contrast optics and a 40X or 60X water-immersion objective (Olympus). Pipettes pulled from thick-walled borosilicate glass capillaries (WPI) had open tip resistances of 3–5 MΩ and 2–3 MΩ for the pre- and postsynaptic recordings, respectively. Whole-cell current- and voltage-clamp recordings were made with a Multiclamp 700B amplifier (Molecular Devices, Foster City, CA). Calyx terminals and MNTB neurons were identified visually by their appearance in contrast optics and/or presynaptic fluorescence of Alexa 594. For presynaptic rat calyx of Held recordings, pipettes contained (in mM): 110 K-gluconate, 20 KCl, 1 MgCl_2_, 10 HEPES, 4 MgATP, 0.01-0.02 Alexa 594, 0.3 Tris-GTP, and 3 Na_2_-phosphocreatine and 10 Tris_2_-phosphocreatine (290 mOsm; pH 7.3 with KOH). For postsynaptic rat MNTB principal neuron recording, pipettes contained (in mM) 130 CsCl, 10 HEPES, 9 NaCl, 10 EGTA, 1 MgCl_2_, 5 disodium phosphocreatine, 2 QX-314, 4 MgATP, 0.3 Na_2_GTP, pH 7.3 at 295 mOsm, pH 7.3 with CsOH. For postsynaptic mouse MNTB, pipettes contained (in mM) 115 TEA-Cl, 4.5 MgCl_2_, 3.5 QX-314-Cl, 10 HEPES, 10 EGTA, 4 ATP-Na_2_, 0.5 GTP-Tris, 5 CsOH, 15 sucrose. This latter solution was developed to improve voltage clamp control at depolarized holding potentials by reducing potassium leak current with a high concentration of TEA. For recordings from DCN cartwheel cells, pipettes contained 128.5 CsCl, 4.8 MgCl_2_, 4 ATP, 0.5 GTP, 14 Tris-phosphocreatine, 0.1 EGTA, and 10 HEPES. Cartwheel cells were identified by previously published criteria (Roberts et al. 2008). Series resistances (6–25 MΩ) were compensated by 60%–80% (bandwidth 3 kHz). Signals were filtered at 10 kHz and sampled at 20 kHz. Antagonists were bath applied or applied by local pressure ejection near the soma.

### Synaptic recordings

MNTB: IPSCs were evoked by voltage pulses (0.1 ms, 10-40V) from a theta glass stimulation pipette positioned near the recorded cell. IPSCs were isolated by blocking excitatory transmission with 10 μM NBQX and 5 μM R-CPP in all experiments. DCN: IPSCs were elicited by voltage clamping the presynaptic cell and delivering brief voltage steps (20–100 µs to −20 or - 30 mV) to evoke an escaping simple spike. Miniature IPSCs were recorded in 1 μM tetrodotoxin (TTX) to block spikes, 10 μM SR95531 to block GABA_A_ receptors, and 10 µM NBQX + 5 µM R-CPP (or 50 µM D-AP5) to block glutamate receptors. Individual mIPSCs were detected using a template matching algorithm (AxoGraph X; rise time of 0.2 ms and decay of 0.5 ms, threshold of 3× noise SD) with a minimum amplitude cutoff between −10 and −20 pA. Events were inspected visually for false positives or malformed events.

### Analysis

Data were analyzed using Clampfit (Molecular Devices) and Igor (WaveMetrics). Reported voltages were corrected for junction potentials as indicated. Statistical measurements are expressed as mean ± SEM unless otherwise stated.

## Results

### Effect of XE991 on glycinergic IPSCs

These experiments were initiated during a study of presynaptic Kv7 channels and their role in synaptic transmission (Huang and Trussell, 2011). The first set of recordings were made from principal neurons of the rat MNTB in which presynaptic glycinergic fibers were activated with an extracellular stimulus electrode (Awatramani *et al*., 2004). These neurons express heteromeric alpha1/beta glycine receptor subunits at sites postsynaptic to glycinergic terminals (Awatramani et al., 2004; Hruskova et al. 2012). Postsynaptic cells were recorded with elevated intracellular Cl^-^ to increase the amplitude of the IPSCs recorded while voltage clamping cells at −60 mV. Bath application of 10 μM XE991 reduced the amplitude of the IPSCs to 38.7±5.4 % of control (N=6) (Figure 1A, B). Some evidence for partial reversal of the effect was seen after 10-20 minutes of washout, although experiments did not last long enough to attain complete reversal in most cases. This inhibition of glycinergic IPSCs was unexpected as XE991 was previously shown to increase the amplitude of excitatory synaptic currents onto these same neurons by a presynaptic mechanism (Huang and Trussell, 2011). We therefore examined the paired pulse ratio (PPR) of IPSCs, stimuli at 50-ms intervals, and found no significant effect of the drug on PPR (Fig 1B, C; control ratio 1.00±0.23; XE991 ratio 0.95±0.14, N=8, p=0.59, paired t-test), suggesting a postsynaptic action of XE991.

**Fig. 1.**
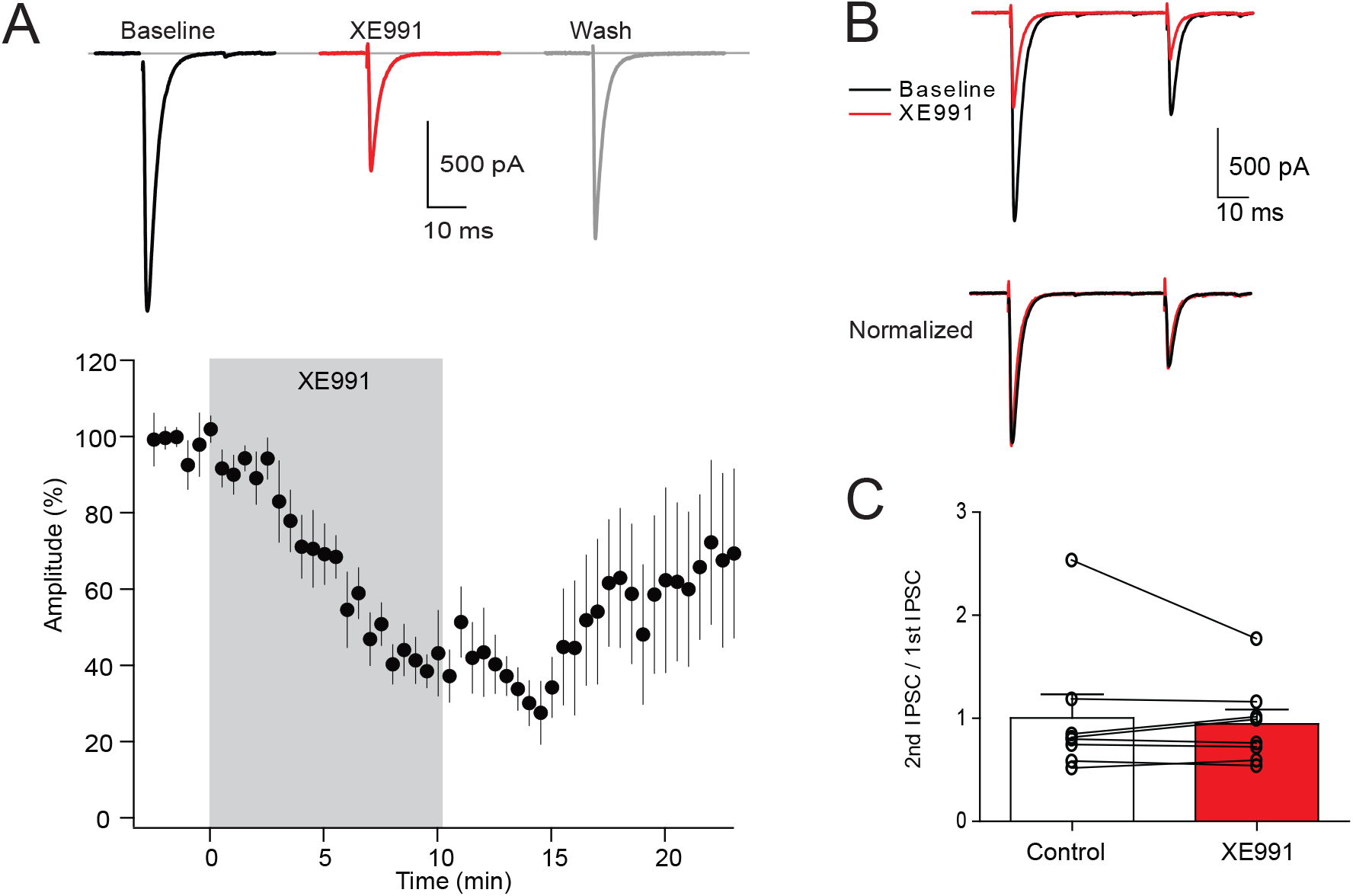
Block of IPSCs in the rat MNTB. A, Top, Block of evoked IPSC with 10 μM XE991 (red trace), and partial reversal. Bottom, time course of action of 10 μM XE991 on evoked IPSCs for 6 cells. Peak amplitudes reduced to 38.7±5.4% of control after 10 min. incubation. Drug effect washed out slowly (more than 10 minutes to return to baseline). B, Paired-pulse protocol (50 ms interval). Traces below are normalized to amplitude of first response to illustrate lack of effect on paired pulse ratio. C, Calculated paired-pulse ratio was not changed after XE991 application (p = 0.59, paired t-test, n=8), indicating XE991 acts on postsynaptic sites.

Given that Kv7 channels may maintain presynaptic resting potential, it was possible that XE991 inhibited the IPSCs by depolarizing inhibitory axons, thereby inactivating Na^+^ channels and elevating spike threshold in axons, and thus reducing the number of axons successfully stimulated on each trial. To eliminate this possibility, paired recordings were established between synaptically coupled cartwheel interneurons in the DCN, in which action potentials in the presynaptic cell reliably elicited IPSCs in the postsynaptic cell on each trial. After establishing a stable baseline of activity, 10 μM XE991 was washed into the bath. The amplitude of IPSCs were reduced over a 10-15 minute period to a stable value of 36.6±0.3% of control (P<0.001, one-sample t-test, n=4 cells) (Fig. 2A-C); during this time there was no systematic change in the paired pulse ratio). In no case did the presynaptic cell fail to elicit a spike. These results again suggest a postsynaptic target of the drug.

**Fig. 2.**
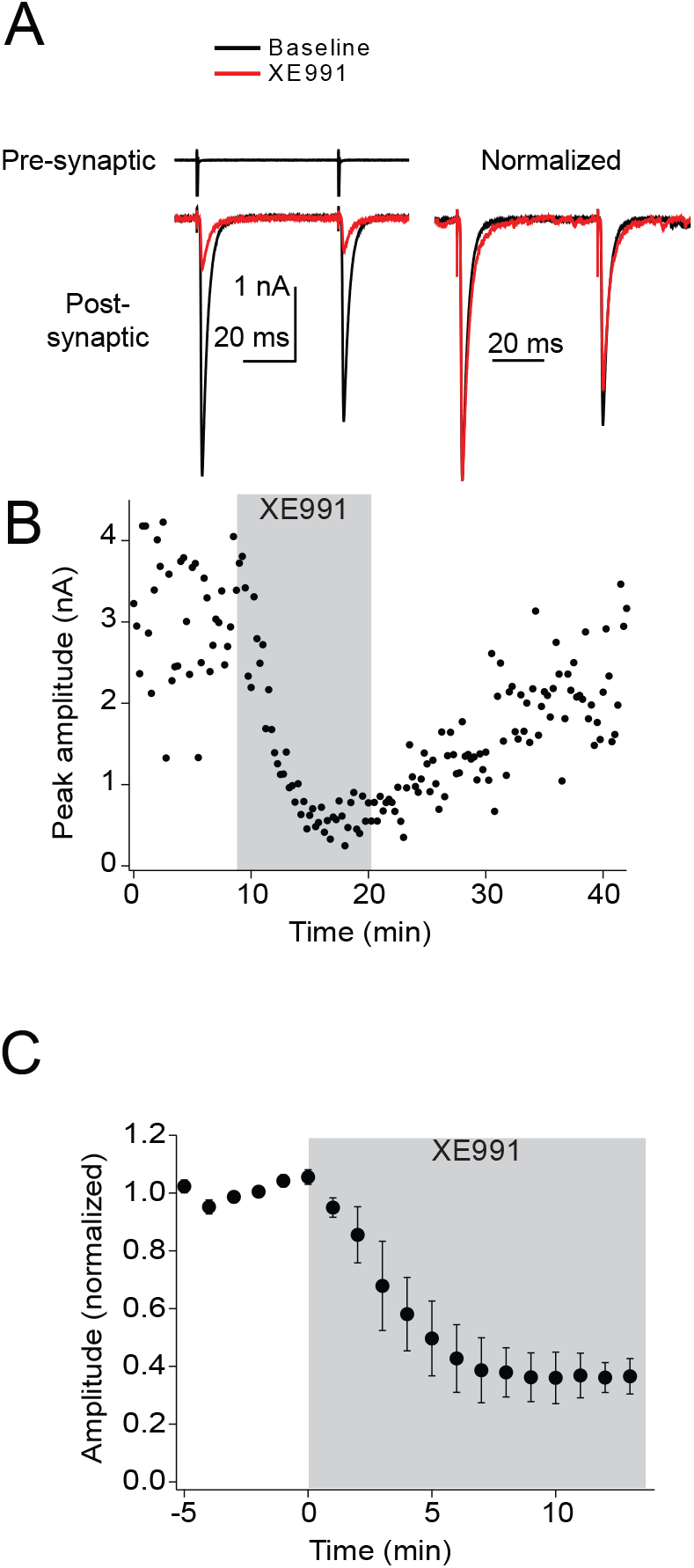
Block of IPSCs in the mouse DCN. A. Two escaping spikes, 50-ms interval, were elicited in presynaptic neuron resulting in postsynaptic IPSCs. Red trace recorded 10 min after wash-in of 10 μM XE991. Left traces are normalized to amplitude of first IPSC in pair. B, diary plot for experiment in panel A shows partial reversal after 20 minutes. C. data averaged from 3 paired and one autaptic recording. Responses were pooled for each 1 minute of recording.

Further evidence for a postsynaptic site of action came from analysis of miniature glycinergic IPSCs (mIPSCs). Miniature IPSCs were recorded in cartwheel cells in the presence of 1 μM TTX to block spikes, and then again with addition of 10 μM XE991 (n=4 cells) or further addition of 100 μM XE991 (n=3 cells; Fig. 3A). Amplitudes of individual events were broadly distributed (Fig 3B), but it was clear that XE911 had a significant dose-dependent reduction in the mean amplitude of mIPSCs (10 μM, 66.0±0.1% of control; 100 μM, 39±0.1% of control; P<0.0001, t-test between 10 μM and 100 μM; Fig. 3A-C), indicating that the drug was acting on postsynaptic glycine receptors.

**Fig. 3.**
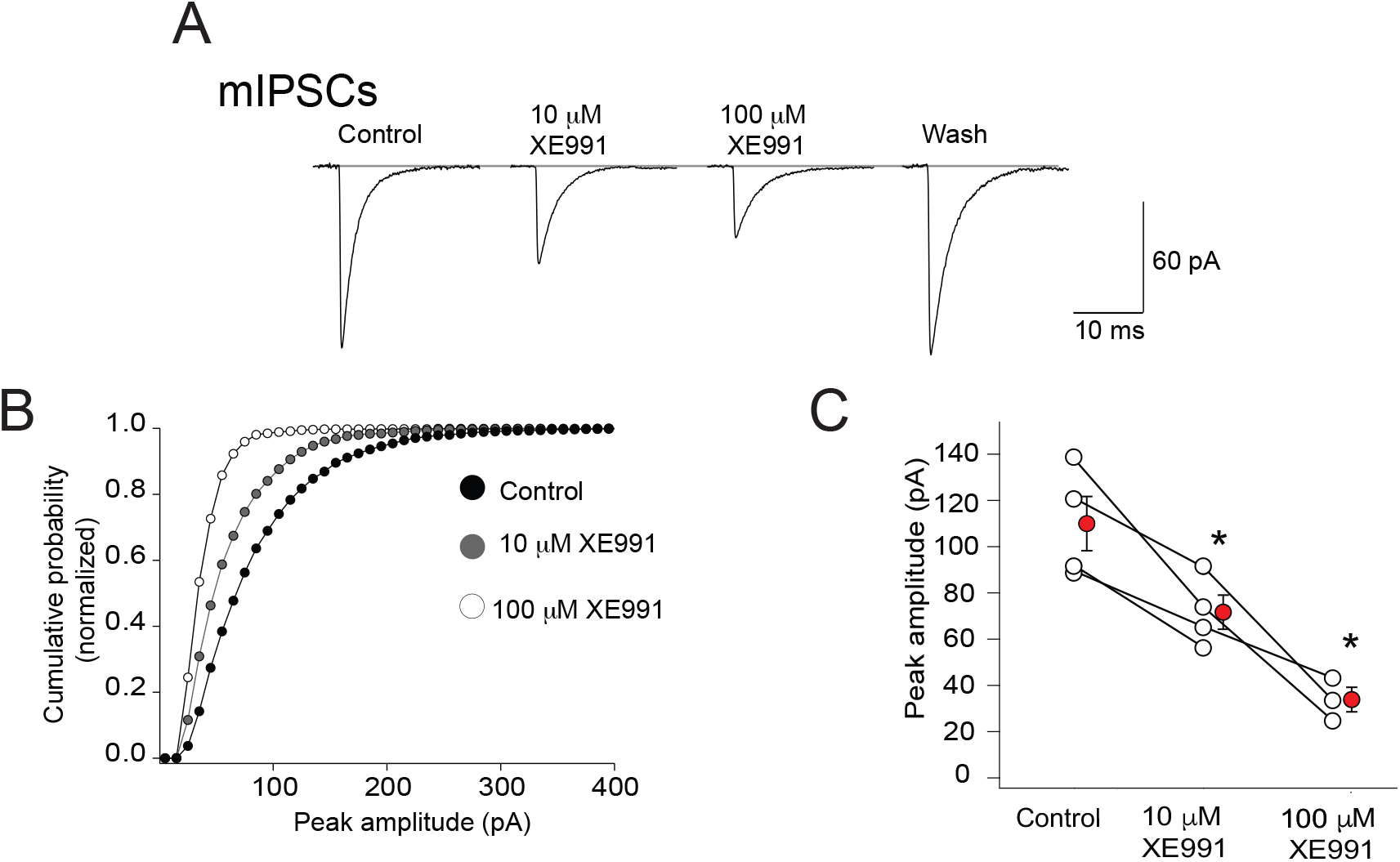
XE991 blocks mIPSCs in mouse DCN. A, Average traces of glycinergic mIPSCs recorded from DCN cartwheel cells. XE991 blocked the peak amplitudes in a dose dependent manner, and the effect was reversible. B, Cumulative probability plots showing leftward shifts of the distribution of mIPSC peak amplitudes in the presence of 10 and 100 μM XE991, suggesting the drug acted at postsynaptic sites. C, Absolute mean peak amplitude of mIPSCs were reduced by XE991 in 4 cells. Lines connect responses from the same recording. Means in red. * Unpaired t test on means: comparison of control to 10 μM: P=0.033; 10 μM to 100 μM, P=0.012.

### Effects of XE991 on responses to exogenous glycine

To test direct effects of XE991 on glycine receptors we made recordings from rat or mouse MNTB and applied glycine to cell bodies by pressure ejection from a nearby pipette. Brief pulses (5-7 ms) of a high concentration of glycine (1 mM) assured that reproducible responses could be obtained while not appreciably diluting the concentration of bath applied drugs near the cell surface. In rat MNTB neurons, XE991 produced significant dose-dependent reduction of the glycine response (10 μM: 49.6±4.4% of control, n = 6; 100 μM, 12.2±1.9% of control, n=6; p<0.001 one-sample t-test; Fig. 4A, B). Notably, these concentrations had no effect on Cl^-^ currents elicited by puffs of 1 mM GABA (Fig 4C; 10 µM: 97.73±4.1% of control, n=6, p=0.6; 100 µM: 89.43±5.2% of control, n=6, p=0.09; one-sample t-test).

**Fig. 4.**
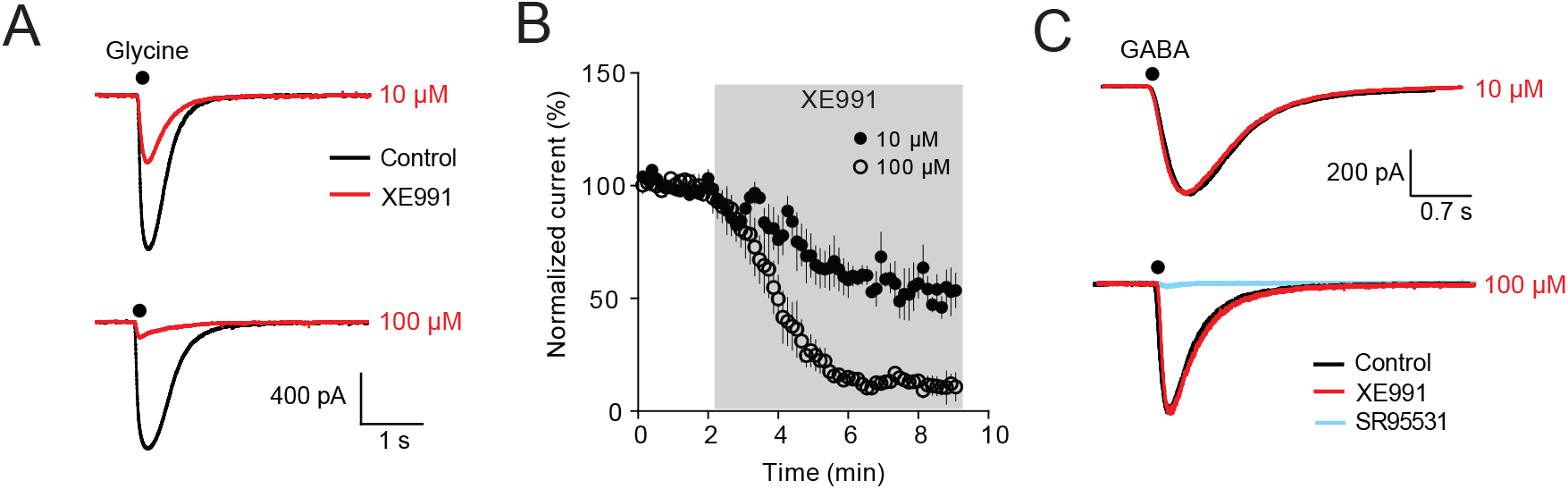
XE991 blocks responses to glycine but not to GABA, in rat MNTB. A. Example traces of effects of 10 μM (top) and 100 μM (bottom) XE991 on responses to puff application of 1 mM glycine to MNTB soma. Drug trace in red. B. Time course of drug effects on puff-evoked glycine responses in MNTB. 10 μM of XE991 reduced the response to 49.6±4.4% of control, n = 6, while 100 μM inhibited the responses to 12.2±1.9% of control, n = 6. C, Neither 10 μM (97.7±4.1% of baseline, p = 0.60, n = 6, one-sample t-test) nor 100 μM (89.4±5.2% of baseline, p = 0.09, n = 6, one-sample t-test) XE991 blocks responses evoked by puff application of 1 mM GABA to MNTB neurons. XE991 trace in red. In the lower panel 5 μM GABA_A_ receptor antagonist SR95531 completely blocked the GABA response (blue trace).

A second series of experiments were conducted in mouse MNTB to further explore the pharmacology and voltage sensitivity of block. In order to better control voltage at depolarized potentials, cells were loaded with a TEA-based solution instead of Cs^+^, and current amplitudes were limited by holding at −30 mV. 100 μM XE991 blocked the response to 1 mM glycine to 32.3±0.1% of control (n=6), somewhat less than the block observed in rat. To determine if the block by XE991 was competitive, the effect was compared for puffs of either 1 mM or 10 mM glycine. While the absolute concentration of glycine on the cell body is uncertain with this method of application, by comparing 1 vs 10 mM puff pipette concentrations, we achieved a ten-fold range of final concentration at the cell surface. The degree of block by 100 μM XE991 was identical in the two cases (10 mM glycine: 33.7±0.1% of control, n=4; p=0.98 compared to 1 mM glycine data above). Thus, XE991 is not likely to be competitive with glycine.

As XE991 is cationic, it is not likely to block the glycine-gated anion pore. Such pore block would be expected to bestow a voltage sensitivity to the glycine conductance. To test this possibility, glycine responses were obtained over a wide range of holding potentials with and without XE991. Fig 5A-B shows represented responses and their current-voltage relation. Profound and reversible block by 100 μM XE991 was obtained at all potentials. Fig. 5C shows data averaged across 6 cells and illustrates that glycine currents were linearly related to holding potential even in the presence of XE991. This linearity is emphasized in Fig 5D in which control and drug data are normalized to the peak current at +42.3 mV, showing that the degree of block was independent of holding potential. We conclude that XE991 generates a non-voltage-dependent, non-competitive block of glycine receptors.

**Fig. 5.**
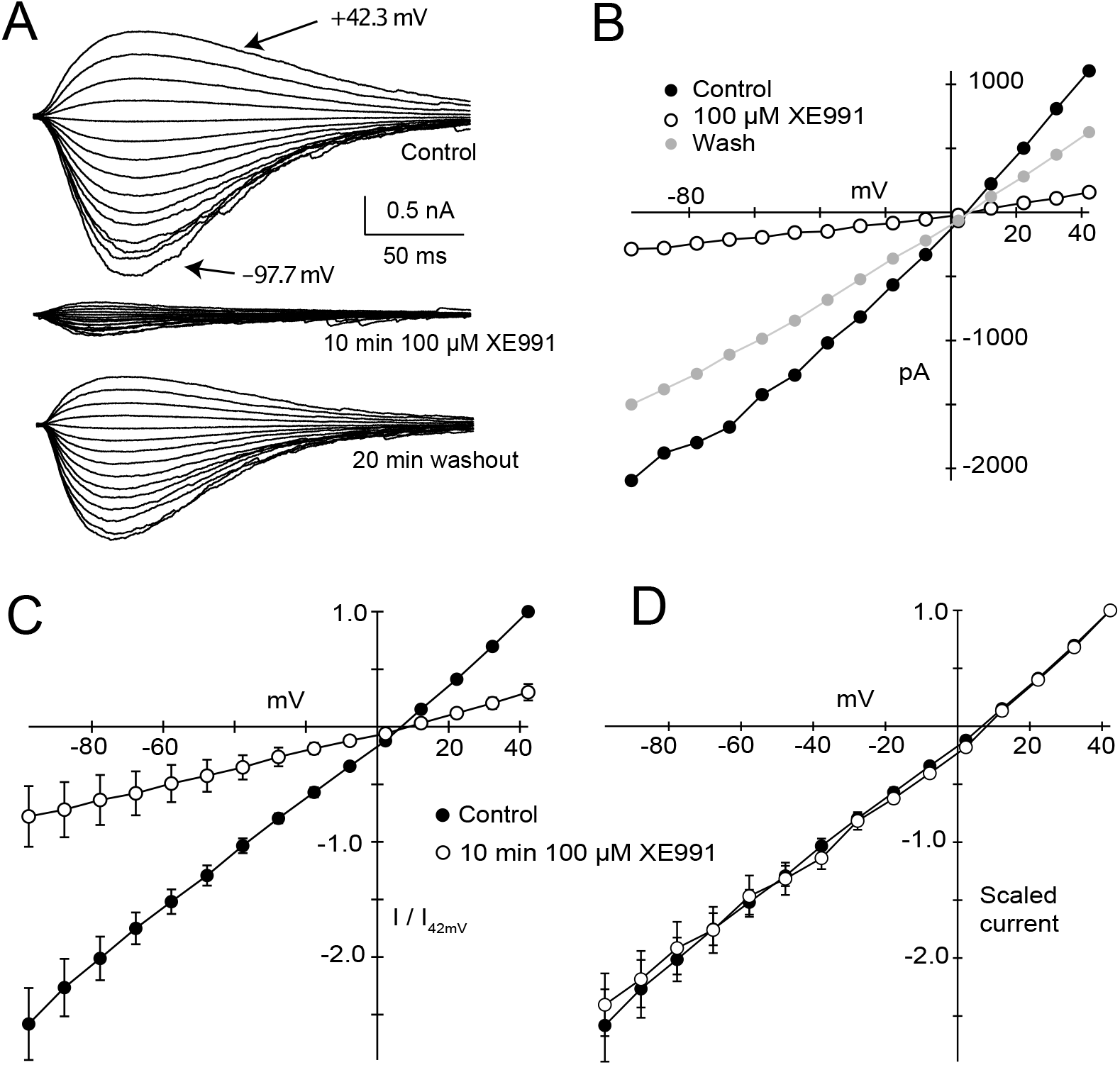
Block of glycine response by XE991 is not voltage dependent. A, Example traces of puff-evoked glycinergic currents at different holding potentials (10 mV intervals) in mouse MNTB neurons. Recordings were made in the control bath solution (top) or in the presence of 100 μM XE991 (middle), and block was almost completely reversible (bottom). B, Current-voltage relationship of glycinergic responses shown in A. XE991 blocks the response at every voltage. C, Current voltage relations averaged across 6 cells, normalized to the control response at +42.3 mV. D, data from C, but each condition was normalized to the +42.3 mV data point, showing that the shape of the curves and reversal potential are unchanged by XE991.

We also used mouse MNTB to examine two other compounds used routinely in studies of Kv7 channels. Linopirdine appeared somewhat more potent than XE991, inhibiting the response to 1 mM glycine to 10±4% of control at 50 μM (n=6, p=0.001, one-sample t-test). By contrast, no effect on the glycine response was observed after 10 minutes incubation with the Kv7 channel activator retigabine (10 μM; n=4, P=0.49).

Lastly, we returned to potential presynaptic effects of XE991 by considering whether the drug can also block presynaptic glycine receptors (Fig. 6A, B). Glycine receptors expressed on the calyx of Held, a glutamatergic axon terminal, were shown to contain homomeric alpha 1 glycine receptor subunits, with consequent effects on agonist and antagonist sensitivity (Hruskova *et al*., 2012). Presynaptic whole cell recordings were made from calyx terminals (Turecek and Trussell, 2001). In the presence of 10 μM XE991, currents activated by 7 ms puffs of 1 mM glycine were 33.3% ± 1.0% of control (n = 6), indicating that XE991 inhibition likely does not depend on the presence of the beta subunit of the receptor.

**Fig 6.**
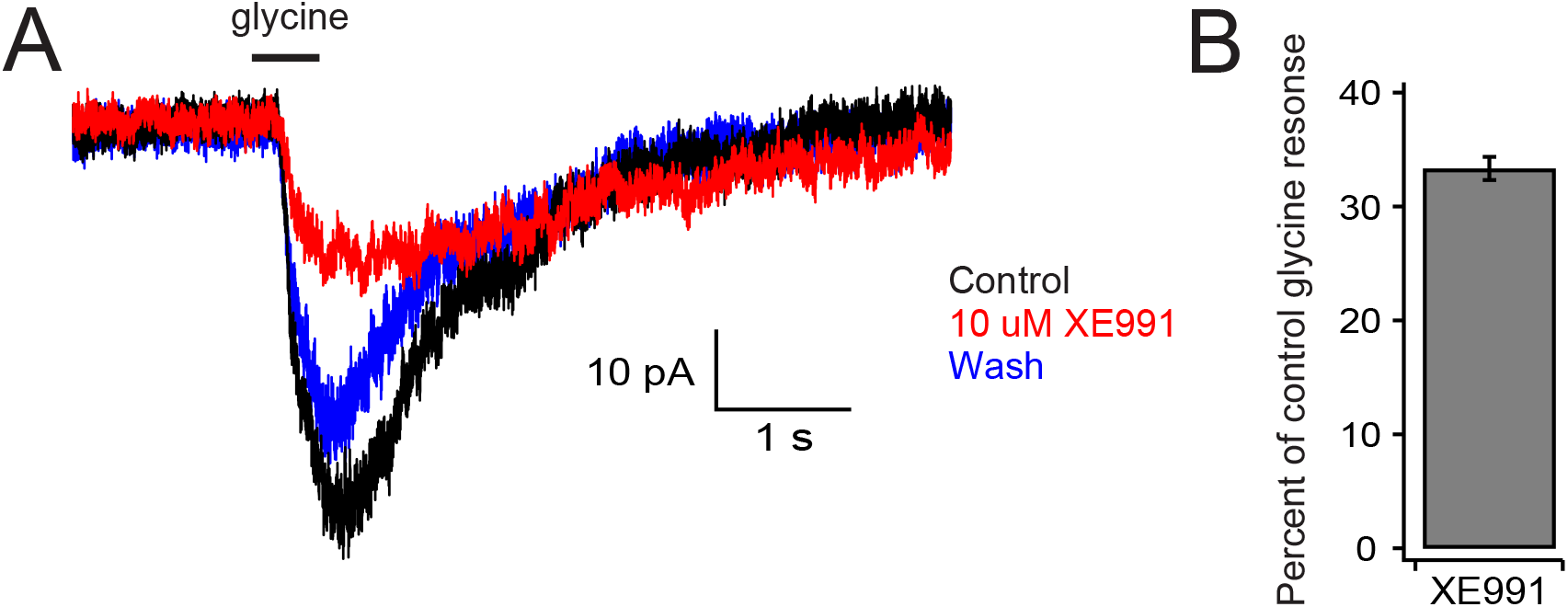
Glycine responses are blocked by XE991 in the rat calyx of Held, a presynaptic nerve terminal. A, example traces showing block by 10 μM XE991 (red) and almost complete reversal (blue). B, group data for fraction of control response in XE991.

## Discussion

This study demonstrates that two common antagonists of Kv7 channels, XE991 and linopirdine, at concentrations typically used in electrophysiological studies, inhibit glycinergic transmission by a postsynaptic mechanism, directly inhibiting activation of glycine receptors. For example, glycine receptors were blocked by ∼50% by 10 μM XE991, while in vitro studies of the drugs actions at Kv7 channels used between three and 40 μM (Huang and Trussell, 2011; Shi *et al*., 2013; Cerina *et al*., 2015; Yamada-Hanff and Bean, 2015; Kuo *et al*., 2016; Peng *et al*., 2017; Martinello *et al*., 2019). By contrast, the IC50 for the drug at KCNQ2/3 channels is ∼0.6 μM (Wang *et al*., 1998). XE991 did not affect GABA_A_ receptors or the shape of the current-voltage relation for glycine responses. Homomeric (presynaptic) or heteromeric (postsynaptic) receptors were blocked similarly. Additionally, XE991 showed equivalent block across a tenfold range in glycine concentrations. We conclude from these observations that the drug inhibits glycine receptors allosterically at the GlyR α1 subunit, but does not directly occlude the glycine-gated chloride channel. At 100 μM, the maximum concentration used, XE991 inhibited glycine receptors by about 68-88% depending on the preparation. However significant inhibition of glycinergic transmission was observed even at 10 μM XE991, a concentration often used in studies of Kv7 channel function. An implication of this study is that investigations using these compounds to explore the roles of Kv7 channels in neural function, at the synaptic or systems level, must account for this potential off-target effect.

Kv7 channels have been shown to control synaptic function in a variety of ways. Somatodendritic Kv7 channels in pyramidal neurons prevent repetitive firing, while axonal Kv7 channels stabilize resting potential, and help restore Na^+^ channel availability and spike duration (Shah *et al*., 2008; Battefeld *et al*., 2014; Hu and Bean, 2018). Such changes in spike duration can regulate calcium influx into nerve terminals. Moreover, presynaptic Kv7 channels set presynaptic resting potential and intracellular Na^+^ concentrations of nerve terminals, both of which impact synaptic strength (Huang and Trussell, 2011). Accordingly, Kv7 channels are proposed as critical components of a wide variety of neural systems. Not surprisingly then, extensive studies have suggested that mutations or modulation in Kv7 may underlie several neurological disorders, such as epilepsy or tinnitus (Shah *et al*., 2008; Grunnet *et al*., 2014; Kalappa *et al*., 2015; Barrese *et al*., 2018), and therefore activators or inhibitors of Kv7 channel are useful probes into disease mechanism and may provide therapeutic strategies.

XE991 was first described as a more potent alternative to linopirdine in studies of hippocampal acetylcholine release (Zaczek *et al*., 1998), with the aim of developing cognitive enhancers for treatment of Alzheimer’s disease. Later studies also utilized XE991 in disease models, for example as a neuroprotective agent in Parkinson’s disease (Liu *et al*., 2018). However, some research has highlighted off-target actions of Kv7 drugs. Linopirdine for example has been shown to antagonize the response of GABA and neuronal nicotinic receptors (Lamas *et al*., 1997), and XE991 may block ERG K+ channels (Elmedyb *et al*., 2007). In this study, XE991 did not inhibit postsynaptic GABA_A_ receptors, consistent with a previous study that showed no effect of the drug on presynaptic GABA_A_ receptors (Huang and Trussell, 2011). Rather, our work focused on an unexpected, potent effect of XE991 on glycine receptors which may impact studies of the functional roles of Kv7 channels in brain. Glycinergic transmission is fundamental to neural circuits of spinal cord, brainstem, cerebellum and midbrain. In these regions, Kv7 channels have been proposed to play significant roles in disease. In the dorsal cochlear nucleus, suppressed Kv7 function is associated with induction of tinnitus following noise exposure, associated with XE991-sensitive hyperactivity of principal cells (Li *et al*., 2013). While these in vitro experiments were performed with glycine receptors blocked, it is likely that in the absence of such blockade, XE991 could further enhance excitability by blocking glycine receptors. Reduced Kv7 expression in sensory neurons is associated with diabetic nerve pain, and here XE991 enhanced mechanical allodynia and thermal hyperalgesia, an effect that could be impacted by inhibition of glycine receptors in spinal circuits. Interestingly, although glycinergic synapses are uncommon in regions higher than midbrain, glycine receptors are found throughout the forebrain, where they are tonically activated by ambient glycine (McCracken *et al*., 2017). In hippocampus, the sensitivity to induction of long-term potentiation is reduced by systemic application of XE991 (Song *et al*., 2009). Moreover, enhanced glycine receptor expression was found in hippocampi of temporal lobe epilepsy patients, suggesting that tonic glycine receptor activity could impact excitability leading to seizure (Eichler *et al*., 2008). Exploration of Kv7 channel function in epileptogenesis and other forebrain disorders using channel antagonists will need to distinguish between XE991’s actions on Kv7 and glycine receptors.

## Acknowledgements

This work was supported by NIH grant DC004450.

## Author contributions

Participated in research design: Apostolides, Huang, Lu, Romero, Trussell.

Conducted experiments: Apostolides, Huang, Lu, Romero.

Performed data analysis: Apostolides, Huang, Lu, Romero, Trussell.

Wrote or contributed to the writing of the manuscript: Apostolides, Lu, Romero, Trussell.

## References

Awatramani GB, Turecek R, and Trussell LO (2004) Inhibitory control at a synaptic relay. J Neurosci 24:2643–2647.

Barrese V, Stott JB, and Greenwood IA (2018) KCNQ-Encoded Potassium Channels as Therapeutic Targets. Annu Rev Pharmacol Toxicol 58:625–648.

Battefeld A, Tran BT, Gavrilis J, Cooper EC, and Kole MHP (2014) Heteromeric Kv7.2/7.3 channels differentially regulate action potential initiation and conduction in neocortical myelinated axons. J Neurosci 34:3719–3732.

Beato M, Burzomato V, and Sivilotti LG (2007) The kinetics of inhibition of rat recombinant heteromeric α1β glycine receptors by the low-affinity antagonist SR-95531. The Journal of Physiology 580:171–179.

Borst JGG, and Soria van Hoeve J (2012) The calyx of Held synapse: from model synapse to auditory relay. Annu Rev Physiol 74:199–224.

Brown DA, and Passmore GM (2009) Neural KCNQ (Kv7) channels. Br J Pharmacol 156:1185– 1195.

Cerina M, Szkudlarek HJ, Coulon P, Meuth P, Kanyshkova T, Nguyen XV, Göbel K, Seidenbecher T, Meuth SG, Pape H-C, and Budde T (2015) Thalamic Kv 7 channels: pharmacological properties and activity control during noxious signal processing. Br J Pharmacol 172:3126–3140.

Elmedyb P, Calloe K, Schmitt N, Hansen RS, Grunnet M, and Olesen S-P (2007) Modulation of ERG channels by XE991. Basic Clin Pharmacol Toxicol 100:316–322.

Grunnet M, Strøbæk D, Hougaard C, and Christophersen P (2014) Kv7 channels as targets for anti-epileptic and psychiatric drug-development. European Journal of Pharmacology 726:133–137.

Haick JM, and Byron KL (2016) Novel treatment strategies for smooth muscle disorders: Targeting Kv7 potassium channels. Pharmacol Ther 165:14–25.

Hruskova B, Trojanova J, Kulik A, Kralikova M, Pysanenko K, Bures Z, Syka J, Trussell LO, and Turecek R (2012) Differential distribution of glycine receptor subtypes at the rat calyx of held synapse. J Neurosci 32:17012–17024.

Hu W, and Bean BP (2018) Differential Control of Axonal and Somatic Resting Potential by Voltage-Dependent Conductances in Cortical Layer 5 Pyramidal Neurons. Neuron 97:1315-1326.e3.

Huang H, and Trussell LO (2011) KCNQ5 channels control resting properties and release probability of a synapse. Nat Neurosci 14:840–847.

Jespersen T, Grunnet M, and Olesen S-P (2005) The KCNQ1 potassium channel: from gene to physiological function. Physiology (Bethesda) 20:408–416.

Jonas P, Bischofberger J, and Sandkühler J (1998) Corelease of two fast neurotransmitters at a central synapse. Science 281:419–424.

Kalappa BI, Soh H, Duignan KM, Furuya T, Edwards S, Tzingounis AV, and Tzounopoulos T (2015) Potent KCNQ2/3-specific channel activator suppresses in vivo epileptic activity and prevents the development of tinnitus. J Neurosci 35:8829–8842.

Kuo F-S, Falquetto B, Chen D, Oliveira LM, Takakura AC, and Mulkey DK (2016) In vitro characterization of noradrenergic modulation of chemosensitive neurons in the retrotrapezoid nucleus. J Neurophysiol 116:1024–1035.

Lamas JA, Selyanko AA, and Brown DA (1997) Effects of a cognition-enhancer, linopirdine (DuP 996), on M-type potassium currents (IK(M)) and some other voltage- and ligand-gated membrane currents in rat sympathetic neurons. Eur J Neurosci 9:605–616.

Li P, and Slaughter M (2007) Glycine receptor subunit composition alters the action of GABA antagonists. Vis Neurosci 24:513–521.

Li S, Choi V, and Tzounopoulos T (2013) Pathogenic plasticity of Kv7.2/3 channel activity is essential for the induction of tinnitus. Proc Natl Acad Sci USA 110:9980–9985.

Liu H, Jia L, Chen X, Shi L, and Xie J (2018) The Kv7/KCNQ channel blocker XE991 protects nigral dopaminergic neurons in the 6-hydroxydopamine rat model of Parkinson’s disease. Brain Res Bull 137:132–139.

Martinello K, Giacalone E, Migliore M, Brown DA, and Shah MM (2019) The subthreshold-active KV7 current regulates neurotransmission by limiting spike-induced Ca2+ influx in hippocampal mossy fiber synaptic terminals. Commun Biol 2:145.

McCracken LM, Lowes DC, Salling MC, Carreau-Vollmer C, Odean NN, Blednov YA, Betz H, Harris RA, and Harrison NL (2017) Glycine receptor α3 and α2 subunits mediate tonic and exogenous agonist-induced currents in forebrain. Proc Natl Acad Sci USA 114:E7179–E7186.

Peng H, Bian X-L, Ma F-C, and Wang K-W (2017) Pharmacological modulation of the voltage-gated neuronal Kv7/KCNQ/M-channel alters the intrinsic excitability and synaptic responses of pyramidal neurons in rat prefrontal cortex slices. Acta Pharmacol Sin 38:1248–1256.

Shah MM, Migliore M, Valencia I, Cooper EC, and Brown DA (2008) Functional significance of axonal Kv7 channels in hippocampal pyramidal neurons. Proc Natl Acad Sci USA 105:7869–7874.

Shi L, Bian X, Qu Z, Ma Z, Zhou Y, Wang K, Jiang H, and Xie J (2013) Peptide hormone ghrelin enhances neuronal excitability by inhibition of Kv7/KCNQ channels. Nat Commun 4:1435.

Song M-K, Cui Y-Y, Zhang W-W, Zhu L, Lu Y, and Chen H-Z (2009) The facilitating effect of systemic administration of Kv7/M channel blocker XE991 on LTP induction in the hippocampal CA1 area independent of muscarinic activation. Neuroscience Letters 461:25–29.

Trussell LO, and Oertel D (2018) Mirocircuits of the dorsal cochlear nucleus, in The Mammalian Auditory Pathways: Synaptic Organization and Microcircuits (Oliver DL, Cant NB, Fay RR, and Popper AN eds) pp 73–99, Springer-Verlag.

Turecek R, and Trussell LO (2001) Presynaptic glycine receptors enhance transmitter release at a mammalian central synapse. Nature 411:587–590.

Wang HS, Pan Z, Shi W, Brown BS, Wymore RS, Cohen IS, Dixon JE, and McKinnon D (1998) KCNQ2 and KCNQ3 potassium channel subunits: molecular correlates of the M-channel. Science 282:1890–1893.

Yamada-Hanff J, and Bean BP (2015) Activation of Ih and TTX-sensitive sodium current at subthreshold voltages during CA1 pyramidal neuron firing. J Neurophysiol 114:2376– 2389.

Zaczek R, Chorvat RJ, Saye JA, Pierdomenico ME, Maciag CM, Logue AR, Fisher BN, Rominger DH, and Earl RA (1998) Two new potent neurotransmitter release enhancers, 10,10-bis(4-pyridinylmethyl)-9(10H)-anthracenone and 10,10-bis(2-fluoro-4-pyridinylmethyl)-9(10H)-anthracenone: comparison to linopirdine. J Pharmacol Exp Ther 285:724–730.

